# Monitoring lasting changes to brain tissue integrity through mechanical properties following adolescent exercise intervention in a rat model of Fetal Alcohol Spectrum Disorders

**DOI:** 10.1101/2023.09.26.559571

**Authors:** Katrina A. Milbocker, L. Tyler Williams, Diego A. Caban-Rivera, Ian F. Smith, Samuel Kurtz, Matthew D.J. McGarry, Bertrand Wattrisse, Elijah E.W. Van Houten, Curtis L. Johnson, Anna Y. Klintsova

## Abstract

**Background:** Fetal Alcohol Spectrum Disorders (FASD) encompass a group of highly prevalent conditions resulting from prenatal alcohol exposure. Alcohol exposure during the third trimester of pregnancy overlapping with the brain growth spurt is detrimental to white matter growth and myelination, particularly in the corpus callosum, ultimately affecting tissue integrity in adolescence. Traditional neuroimaging techniques have been essential for assessing neurodevelopment in affected youth; however, these methods are limited in their capacity to track subtle microstructural alterations to white matter, thus restricting their effectiveness in monitoring therapeutic intervention. In this preliminary study we use a highly sensitive and clinically translatable Magnetic Resonance Elastography (MRE) protocol for assessing brain tissue microstructure through its mechanical properties following an exercise intervention in a rat model of FASD.

**Methods:** Rat pups were divided into two groups: alcohol-exposed (AE) pups which received alcohol in milk substitute (5.25 g/kg/day) via intragastric intubation on postnatal days (PD) four through nine during the rat brain growth spurt (Dobbing and Sands, 1979), or sham-intubated (SI) controls. In adolescence, on PD 30, half AE and SI rats were randomly assigned to either a modified home cage with free access to a running wheel or to a new home cage for 12 days (Gursky and Klintsova, 2017). Previous studies conducted in the lab have shown that 12 days of voluntary exercise intervention in adolescence immediately ameliorated callosal myelination in AE rats (Milbocker et al., 2022, 2023). MRE was used to measure longitudinal changes to mechanical properties of the whole brain and the corpus callosum at intervention termination and one-month post-intervention. Histological quantification of precursor and myelinating oligoglia in corpus callosum was performed one-month post-intervention.

**Results:** Prior to intervention, AE rats had lower forebrain stiffness in adolescence compared to SI controls (*p* = 0.02). Exercise intervention immediately mitigated this effect in AE rats, resulting in higher forebrain stiffness post-intervention in adolescence. Similarly, we discovered that forebrain damping ratio was lowest in AE rats in adolescence (*p* < 0.01), irrespective of intervention exposure. One-month post-intervention in adulthood, AE and SI rats exhibited comparable forebrain stiffness and damping ratio (p > 0.05). Taken together, these MRE data suggest that adolescent exercise intervention supports neurodevelopmental “catch-up” in AE rats. Analysis of the stiffness and damping ratio of the body of corpus callosum revealed that these measures increased with age. Finally, histological quantification of myelinating oligodendrocytes one-month post-intervention revealed a negative rebound effect of exercise cessation on the total estimate of these cells in the body of corpus callosum, irrespective of treatment group which was not convergent with noninvasive MRE measures.

**Conclusions:** This is the first application of MRE to measure changes in brain mechanical properties in a rodent model of FASD. MRE successfully captured alcohol-related changes to forebrain stiffness and damping ratio in adolescence. These preliminary findings expand upon results from previous studies which used traditional diffusion neuroimaging to identify structural changes to the adolescent brain in rodent models of FASD (Milbocker et al., 2022; Newville et al., 2017). Additionally, *in vivo* MRE identified an exercise-related alteration to forebrain stiffness that occurred in adolescence, immediately post-intervention.

## INTRODUCTION

1 in 20 infants are diagnosed with Fetal Alcohol Spectrum Disorders (FASD) in the United States annually (Hoyme et al., 2016; Popova et al., 2018). Despite the global prevalence of FASDs, there is a paucity of methods to accurately and reliably diagnose and track FASD progression, impeding development and assessment of effective interventions aimed at improving brain health in this population. Magnetic resonance imaging is a powerful noninvasive tool that has been used to identify FASD-specific anomalies in brain structure and function across the lifespan (Alger et al., 2021; Lebel and Ware, 2023; Lewis et al., 2021; Norman et al., 2009; O’Neill et al., 2019; Rockhold et al., 2023). For instance, MRI studies have revealed that corpus callosum development is highly vulnerable to alcohol teratogenesis and is dependent on the timing and amount of alcohol exposure. Deficits in callosal growth and organization during adolescence have been correlated with delayed math/language capacities, impaired bimanual coordination and disrupted visuospatial processing in youth with FASD (Gimbel et al., 2022; Mattson et al., 2019; Roebuck-Spencer and Mattson, 2004; Sowell et al., 2008; Wozniak et al., 2009).

Based on these findings, targeted therapies that support callosal development in youth with FASD are of significant interest, but none have yet been translated to clinical use. Increasing aerobic activity has been proposed as an accessible behavioral intervention for youth with FASD as exercise upregulates myelination of callosal axons in the typically-developing brain (reviewed in Loprinzi et al., 2020; Pritchard Orr et al., 2018). Indeed, several preclinical studies have demonstrated that callosal maturation is supported by aerobic activity in rodent models of FASD (reviewed in Klintsova et al., 2013). Unfortunately, noninvasive imaging studies examining such responses to aerobic activity are limited as few methods are available to reliably capture short-term changes to brain microstructure. The innovation of novel neuroimaging techniques to track subtle changes to brain microstructure longitudinally are imperative for tailoring exercise-related intervention strategies for FASD affected youth.

Magnetic resonance elastography (MRE) offers a quantitative MRI contrast sensitive to brain tissue microstructure through its viscoelastic mechanical properties (Arani et al., 2021; Hiscox et al., 2016; Murphy et al., 2019). Two mechanical properties are commonly reported outcomes of MRE scanning: tissue stiffness and damping ratio. In the adult brain, higher global shear stiffness corresponds to healthier tissue, and stiffness decreases with age (reviewed in Hiscox et al., 2021) and in neurodegenerative conditions including multiple sclerosis (Streitberger et al., 2012; Wuerfel et al., 2010) and Alzheimer’s disease (Hiscox et al., 2020a; Murphy et al., 2011). White matter is stiffer than gray matter in healthy adults (Johnson et al., 2013b; van Dommelen et al., 2010), and within white matter highly organized tracts like corpus callosum are stiffest (Smith et al., 2022a). Damping ratio, a dimensionless parameter that describes motion attenuation due to energy dissipation, is lower in highly organized white matter tracts like corpus callosum (Johnson et al., 2013a). Damping ratio is less commonly reported in the MRE literature but has notably been associated with cognitive function in healthy adults (Delgorio et al., 2022; Hiscox et al., 2020b).

Similar findings are described in the preclinical MRE literature, and where, the application of MRE to rodent models of neurodevelopmental disorders is expanding. Global brain stiffness is reduced in models of Alzheimer’s Disease, multiple sclerosis, stroke and in the minutes following respiratory arrest or hypothermic conditions (Bertalan et al., 2020; Freimann et al., 2013; Guo et al., 2019; Millward et al., 2015; Munder et al., 2018). Global brain stiffness increases with age, with pubertal onset in rats and mice, and has been associated with increased myelination and cytoskeletal linking (Guo et al., 2019; Pong et al., 2016). Liu et al. (2023) discovered that *cortical* stiffness increased between PD 28 (juvenile) and 70 (young adult) in female Sprague Dawley rats (Liu et al., 2023). However, mechanical property ranges and trends vary due to differences in scan acquisition, scanner strength, inversion algorithm, and age of the rodents. This pilot study is the first to collect longitudinal brain MRE data in a rat model of FASD. The application of MRE to the investigation of neurological diseases augments our capacity to monitor disease progression and supplements data on microstructural alterations gleaned from conventional neuroimaging methods.

This pilot study is the first to employ a novel MRE protocol using nonlinear inversion (NLI) to characterize microstructural changes in forebrain and corpus callosum after adolescent exercise intervention in a rat model of FASD. We hypothesized that exercise intervention would accelerate forebrain and corpus callosum stiffening in FASD rats, resulting in neurodevelopmental “catch up.” Conversely, we predicted that global and callosal damping ratio would be highest in FASD rats without intervention exposure, indicating higher energy attenuation, and that aerobic activity would normalize tissue damping in FASD rats. Finally, we expected that such changes would be reflected in alterations to the number of precursor and myelinating oligodendrocytes of the corpus callosum as variations to the oligodendrocyte matrix have been implicated in changes to tissue stiffness and damping ratio.

## MATERIALS AND METHODS

**Figure 1** illustrates the experimental timeline for this study including a model of alcohol exposure, exercise intervention, MRI scanning, and immunohistochemistry. All procedures were performed in accordance with an approved University of Delaware IACUC protocol and followed NIH ARRIVE guidelines for animal care.

**Figure 1.**
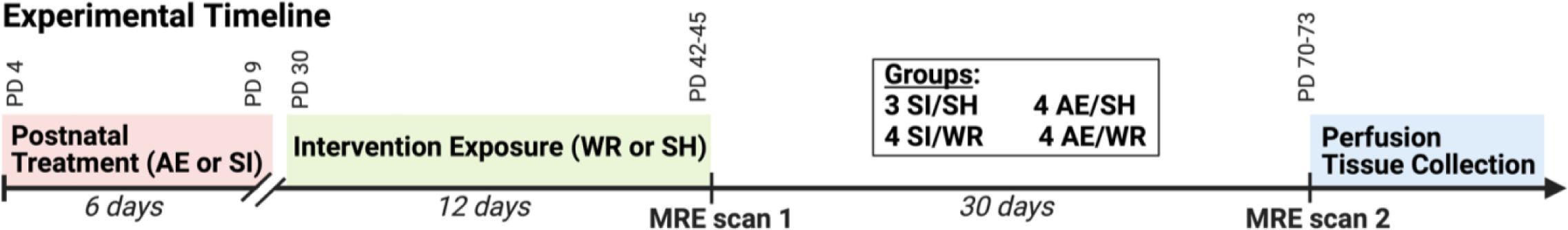
Experimental timeline depicting AE during the brain growth spurt, adolescent voluntary exercise intervention, and noninvasive longitudinal neuroimaging prior to humane euthanasia in young adulthood.

### A. Rat Model of Fetal Alcohol Spectrum Disorders (FASD)

A total of 16 female Long-Evans rat pups from four separate litters were generated for this study and divided evenly between alcohol-exposed (AE) and sham procedural control (SI) groups. A comprehensive protocol was used to model AE during the third trimester of pregnancy targeting the rat brain growth spurt, as described previously (Gursky and Klintsova, 2017; Milbocker and Klintsova, 2020). AE rats were administered 5.25 g/kg/day of ethanol in milk substitute via intragastric intubation twice daily during the rat brain growth spurt on postnatal days (PDs) four through nine (Dobbing and Sands, 1979) yielding a mean BAC of between 300-400 ml/dl. To prevent nutrient loss due to insufficient suckling during intoxication, rat pups in the AE group received two milk-only doses two and four hours after second alcohol dose on PD 4 and one milk-only dose on PD 5-9. SI control rat pups received intragastric intubation without liquid administration to control for the stress of continual intubation during the brain growth spurt (**Figure 1**). Pups were weaned on PD23 and housed in same-sex pairs or triplicates until the start of the intervention period.

### B. Voluntary exercise intervention in adolescence

One week following weaning (PD 30), AE and SI rats were randomly selected to be placed into modified wheel running cages (WR) or new home cages in pairs or triplicates (socially-housed, SH). The final sizes for each of the four groups, comprising both alcohol and intervention conditions, were: AE/WR = 4, AE/SH = 4, SI/WR = 4, and SI/SH = 3. MRE data for one SI/SH rat was discarded as it fell outside of physiological limits likely due to poor scan quality. Histology data was also not collected for this rat. In the modified WR cages, rats had free access to metal running wheels (Lafayette Instrument Company, Lafayette, IN, United States) for 24 hours/day from PD 30-42 and the number of rotations per cage was recorded daily at 9:00 AM (Gursky and Klintsova, 2017). On average, the eight WR rats ran about 115,675 meters total over the 12 days of voluntary wheel running.

### C. MRE Scanning

Rats were MRE scanned twice: once immediately after intervention termination (between PD 42-45) and approximately 30 days following intervention termination (PD 70-73) given scanner availability. All imaging was performed on a Bruker 9.4T Biospec preclinical MRI scanner (Bruker Corporation, Billerica, MA) using a four-channel surface coil. Rats were anesthetized with 1-3% isoflurane administered via nose cone. Respiration and body temperature were monitored throughout the scan session, and temperature was regulated by a liquid heated animal bed. For the MRE scans, 800 Hz vibrations were applied using a nonmagnetic, piezoelectric actuator (APA150M-NM; Cedrat Technologies, Meylan, France). A custom holder and adjustable bite bar transmitted waves from the actuator to the rodent’s brain with amplitudes on the order of 10 μm. A custom MRE spin-echo, echoplanar imaging (EPI) sequence with motion encoding gradients (MEGs) applied in three directions, with four phase offsets, and positive and negative gradient polarities, was used to capture 3D wave motion for a total of 24 repetitions per image. MEGs were unilateral and frequency matched to the actuator vibration, with a train of 19 MEGs comprising the entire encoding block. Magnitude and phase images were acquired using the following imaging parameters: TE/TR = 60/3400 ms, FOV = 20 x 20 mm2, matrix = 80 x 80, slice thickness = 0.5 mm, slices = 40, averages = 24. A resolution of 0.25 x 0.25 x 0.5 mm3 was achieved in 32 minutes. Total scan time including other anatomical images was approximately one hour.

Mechanical properties were estimated from displacement data using the nonlinear inversion (NLI) with a no boundary condition formulation (Kurtz et al., 2022; McGarry et al., 2012), which iteratively solves the Navier equation inverse problem via a finite element mesh and subzone estimation scheme. NLI outputs storage modulus (*G*’) and loss modulus (*G’’*) which are the real and imaginary components of complex shear modulus (*G* = *G*′ + *G’’*). These parameters were then used to calculate shear stiffness (μ=2|*G*|2/(*G*′+|*G*|)) and damping ratio (ξ=*G*′′/2*G*′) (Manduca et al., 2021). NLI parameters originally optimized for the human brain were adjusted to the rat brain based on the imaging volume size, resolution, vibration frequency, and approximate wavelength (Williams et al., 2022): initial property guesses of *G*0’ = 5 kPa and *G*0*’’* = 1 kPa; subzone size = 3 x 3 x 3 mm^3^; spatial filtering width = 0.18 mm. NLI returns voxel-wise material property maps and ROI-based analyses were conducted on the global forebrain and body of the corpus callosum.

### D. Histological quantification of oligoglia one month post-intervention

Rats were humanely euthanized in adulthood following the second scanning session (PD 70-73, approximately 30 days post-intervention) via non-lethal intraperitoneal injection of 150 mg/kg ketamine-xylazine followed by transcardial perfusion with 100 mL heparinized (5000 units/1L solution) phosphate buffer solution (PBS, pH = 7.4) followed by 100 mL of 4% paraformaldehyde in PBS. Brain tissue was collected and post-fixed in 4% paraformaldehyde in PBS that was swapped with a 30% sucrose in 4% paraformaldehyde in PBS mixture three times prior to sectioning. The cerebellum and olfactory bulbs were removed, and the forebrain of each rat was sectioned at a thickness of 40 μm. Sections were stored in rostrocaudal order and anatomical organization was maintained during staining of free-floating sections.

To visualize mature oligodendrocytes (OLs) and oligodendrocyte precursor cells (OPCs), immunocytochemical labeling was performed following procedures described in Milbocker et al. (2023). Unbiased stereological estimation of each cell type was performed through the rostrocaudal extent of the body of the corpus callosum in every 8th section, requiring quantification from 5-7 tissue sections/rat (grid size = 274 μm × 274 μm; counting frame = 150 μm × 150 μm; region of interest percent quantified = 30%). Coefficient of error values never exceeded 0.1 for total cell estimation of each oligoglia subtype (Slomianka and West, 2005; West et al., 1991). Quantification of OLs and OPCs at the first time point (immediately following intervention exposure) in AE and SI rats was performed in a separate cohort of rats and results were described in Milbocker et al. (2023). This data is included here since the conditions of AE and WR intervention were identical.

### E. Statistical analysis

MRE measures were compared within-rat across time using mixed repeated measures ANOVA with Greenhouse-Geisser correction for lack of sphericity. To comprehensively evaluate alterations to brain stiffness and damping ratio in adolescence and adulthood between treatment groups, two-way ANOVA were performed with data from each scanning time point. *Post hoc* analysis with Bonferroni correction for multiple comparisons was used to determine specific group differences. Summary statistics and mean property values for both global forebrain and corpus callosum are given in **Tables 1 and 2**.

**Table 1.** Results from factorial and repeated measures ANOVA comparing the effects of postnatal treatment (AE or SI control) and intervention exposure (WR or SH control) on shear stiffness and damping ratio globally and in corpus callosum at 0 and 30 days since intervention (DSI) termination.

| Brain Region | DSI | DV | Predictor | Df | F | $p$ | $\eta^2 p$ | Bonferroni *post hoc* |
| --- | --- | --- | --- | --- | --- | --- | --- | --- |
| Global | 0 | Stiffness | Postnatal treatment x Intervention | 1,11 | 6.17 | 0.03 | 0.36 | $p = 0.02$ |
|  |  | Damping Ratio | Postnatal treatment | 1,11 | 19.85 | < 0.01 | 0.62 |  |
|  | 30 | Stiffness | n.s. |  |  |  |  |  |
|  |  | Damping Ratio | n.s. |  |  |  |  |  |
| | 0-30 | Stiffness | Time x Intervention | 1,11 | 11 | 0.02 | 0.42 | In SH rats only $p = 0.02$ |
| | | Damping Ratio | Time x Postnatal treatment | 1,11 | 18.08 | < 0.01 | 0.62 | In AE rats only $p = 0.02$ |
| Corpus callosum | 0 | Stiffness | n.s. |  |  |  |  |  |
|  |  | Damping Ratio | Postnatal treatment | 1,11 | 8.37 | 0.01 | 0.41 |  |
|  | 30 | Stiffness | n.s. |  |  |  |  |  |
|  |  | Damping Ratio | n.s. |  |  |  |  |  |
|  | 0-30 | Stiffness | Time | 1,11 | 216.28 | < 0.01 | 0.95 |  |
| | | | Time x Intervention | 1,11 | 8.02 | 0.02 | 0.42 | In WR rats only $p < 0.01$ |
|  |  | Damping Ratio | Time | 1,11 | 66.03 | < 0.01 | 0.86 |  |

**Table 2.** Mean and standard deviation of shear stiffness and damping ratio for SI/SH, SI/WR, AE/SH and AE/WR groups at 0 and 30 days since intervention (DSI) termination.

| Group | DSI | n | Shear Stiffness Mean (SD) | Damping Ratio Mean (SD) | Shear Stiffness Mean (SD) | Damping Ratio Mean (SD) |
| --- | --- | --- | --- | --- | --- | --- |
| SI/SH | 0 | 3 | 6.44 (0.16) | 0.29 (0.08) | 3.08 (0.24) | 0.17 (0.19) |
|  | 30 | 3 | 6.77 (0.35) | 0.30 (0.03) | 5.46 (0.95) | 0.33 (0.11) |
| SI/WR | 0 | 4 | 6.54 (0.32) | 0.31 (0.05) | 3.08 (0.35) | 0.19 (0.02) |
|  | 30 | 4 | 6.39 (0.16) | 0.26 (0.02) | 6.22 (0.50) | 0.29 (0.06) |
| AE/SH | 0 | 4 | 5.77 (0.55) | 0.24 (0.05) | 3.42 (1.41) | 0.14 (0.03) |
|  | 30 | 4 | 6.36 (0.31) | 0.30 (0.07) | 5.16 (1.20) | 0.28 (0.06) |
| AE/WR | 0 | 4 | 6.74 (0.10) | 0.20 (0.02) | 2.97 (0.06) | 0.13 (0.02) |
|  | 30 | 4 | 6.47 (0.49) | 0.29 (0.03) | 6.31 (0.80) | 0.29 (0.06) |

## RESULTS

### A. Global changes to brain viscoelastic properties at 0 to 30 days post-intervention

Global mechanical properties were determined as the average stiffness across the entire forebrain volume mask in central slices around the corpus callosum, in both time points: immediately after exercise cessation and one month post-intervention. Analysis of global shear stiffness at time point one using a two-way ANOVA revealed that there was a significant postnatal treatment x intervention interaction (F1 11 = 6.17, *p* = 0.02, ηp^2^ = 0.36; **Figure 2A**) wherein AE/SH rats have the softest forebrain, indicating potential pathological status or underdevelopment. This is expected given that delayed brain maturation is a hallmark characteristic of FASD (Riley et al., 1995; Treit et al., 2013; Wozniak et al., 2009). Notably, global shear stiffness in AE/WR rats was comparable to that in SI/SH and SI/WR rats, indicating that adolescent exercise intervention immediately enhanced tissue integrity in the FASD brain. Similar analysis at time point two in adulthood, 30 days after intervention, indicated that brain stiffness was comparable between all groups (*p* > 0.05, **Figure 2A**). Indeed, mixed models repeated measures ANOVA revealed a significant time x intervention interaction on global shear stiffness; global shear stiffness increased in AE/SH and SI/SH rats over time but did not change in AE/WR and SI/WR rats (F1 11 = 11, *p* = 0.02, ηp^2^ = 0.42; **Figure 2B**). Together, results indicate that adolescent exercise intervention enhances forebrain tissue integrity in the FASD brain, and this is maintained in adulthood. Results are summarized in **Table 1** and **Table 2** and example stiffness maps are shown in **Figure 2C**.

**Figure 2.**
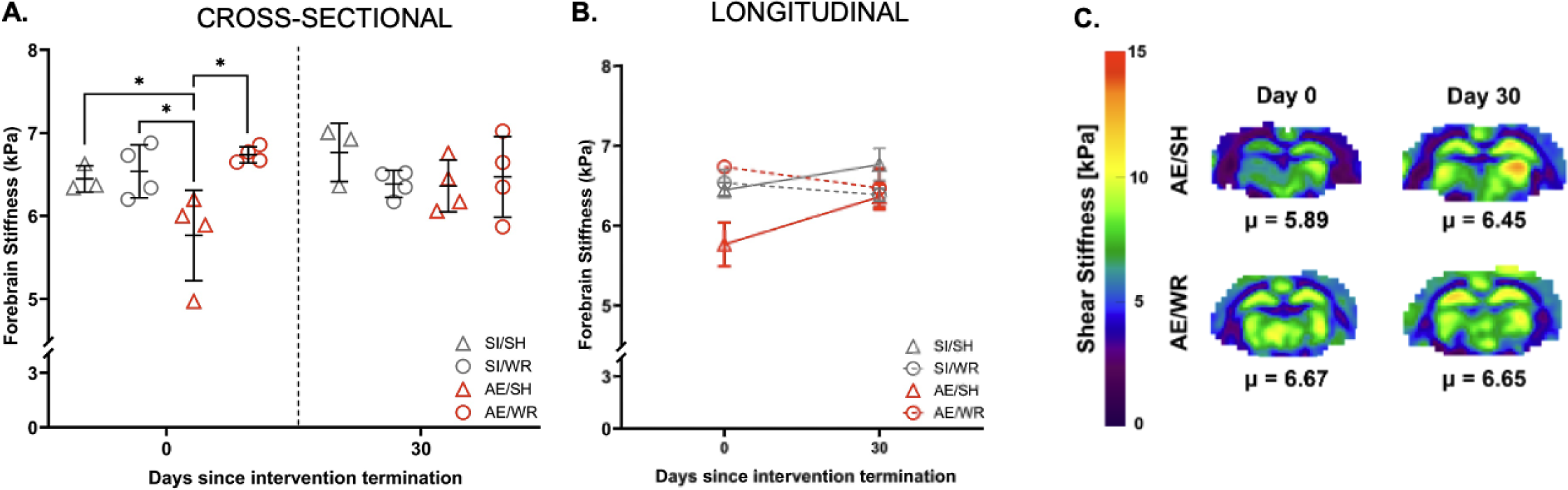
Global shear stiffness measured immediately post-intervention and one month later in adulthood. A) Cross-sectional analysis of global shear stiffness data 0- and 30-days post-intervention for AE/WR, AE/SH, SI/WR and SI/SH groups. * p ≤ 0.05, error bars = SD, B) Longitudinal analysis of mean global shear stiffness values for AE/WR, AE/SH, SI/WR and SI/SH groups; significant main effects and interactions not shown, error bars = ± 1 SE. C) Example shear stiffness maps of AE/SH and AE/WR rats over time. Wheel running immediately ameliorated forebrain stiffness in AE/WR rats.

The same analyses were performed on global damping ratio, calculated in the same fashion as global shear stiffness. Our results show that there is a significant main effect of postnatal treatment condition on damping ratio in adolescence (F1 11 = 19.85, *p* < 0.01, ηp^2^ = 0.62); forebrain damping ratio was lowest in AE/SH and AE/WR (**Figure 3A**). Longitudinal analysis with mixed models repeated ANOVA confirmed that there existed a significant time x postnatal treatment interaction (F1 11 = 18.08, *p* < 0.01, ηp2 = 0.62), with global damping ratio increasing in AE/SH and AE/WR rats between the two scanning time points (**Figure 3B**). In adulthood, one month post-intervention, global damping ratio was comparable between groups. Results are summarized in **Tables 1** and **Table 2** and example damping ratio maps are shown in **Figure 3C**.

**Figure 3.**
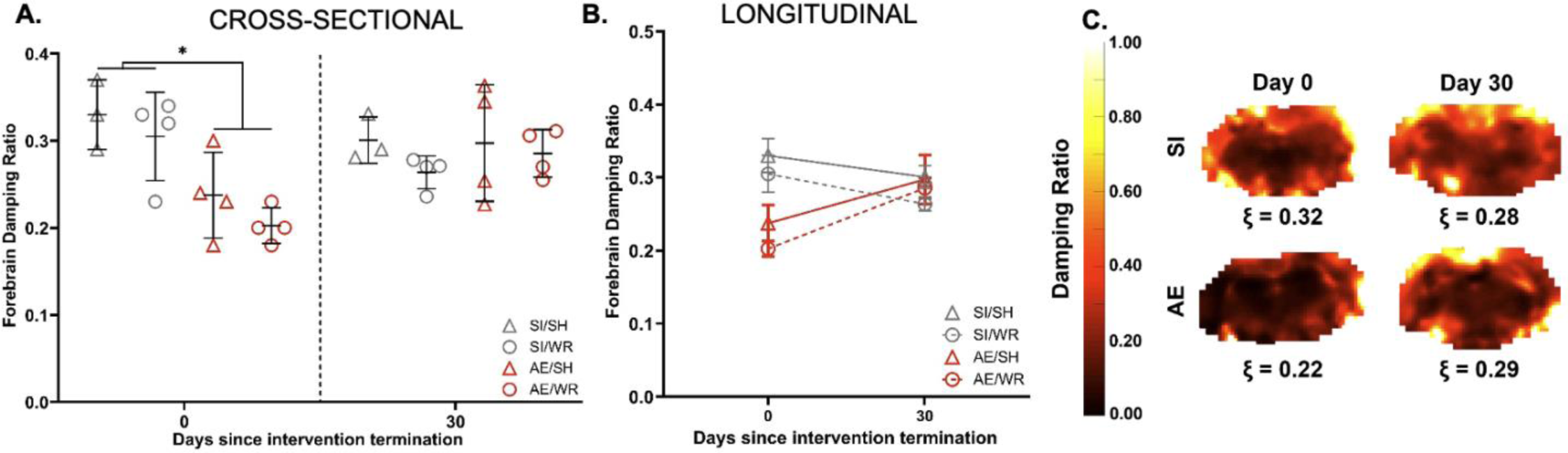
Global damping ratio measured immediately post-intervention and one month later in adulthood. A) Cross-sectional analysis of global damping ratio data 0- and 30-days post-intervention for AE/WR, AE/SH, SI/WR and SI/SH groups. * p ≤ 0.05, error bars = SD, B) Longitudinal analysis of mean global damping ratio values for AE/WR, AE/SH, SI/WR and SI/SH groups. Significant main effects and interactions not shown, error bars = ± 1 SE. AE rats exhibit lower damping ratio than controls 0 days post-intervention, but catch up to control levels 30 days post-intervention. C) Example damping ratio maps of AE and SI rats over time. Damping ratio increased in AE rats from 0- to 30-days post-intervention.

### B. Viscoelastic properties of the body of corpus callosum at 0 to 30 days post-intervention

Regional shear stiffness and damping ratio were calculated in the body of the corpus callosum. Cross-sectional analysis of callosal stiffness using two-way ANOVA demonstrated that shear stiffness did not differ significantly between groups at 0 or 30 days post-intervention (**Figure 4A**). Longitudinal analysis using mixed models repeated measures ANOVA revealed a significant main effect of time (F1 11 = 216.28, *p* < 0.01, ηp^2^ = 0.95) such that all groups were significantly stiffer at time point two. Additionally, there was a significant time x intervention interaction effect wherein the positive change in callosal stiffness was greater in AE/WR and SI/WR rats compared to AE/SH and SI/SH rats (F1 11 = 8.02, *p* = 0.02, ηp2 = 0.42, **Figure 4B**).

**Figure 4.**
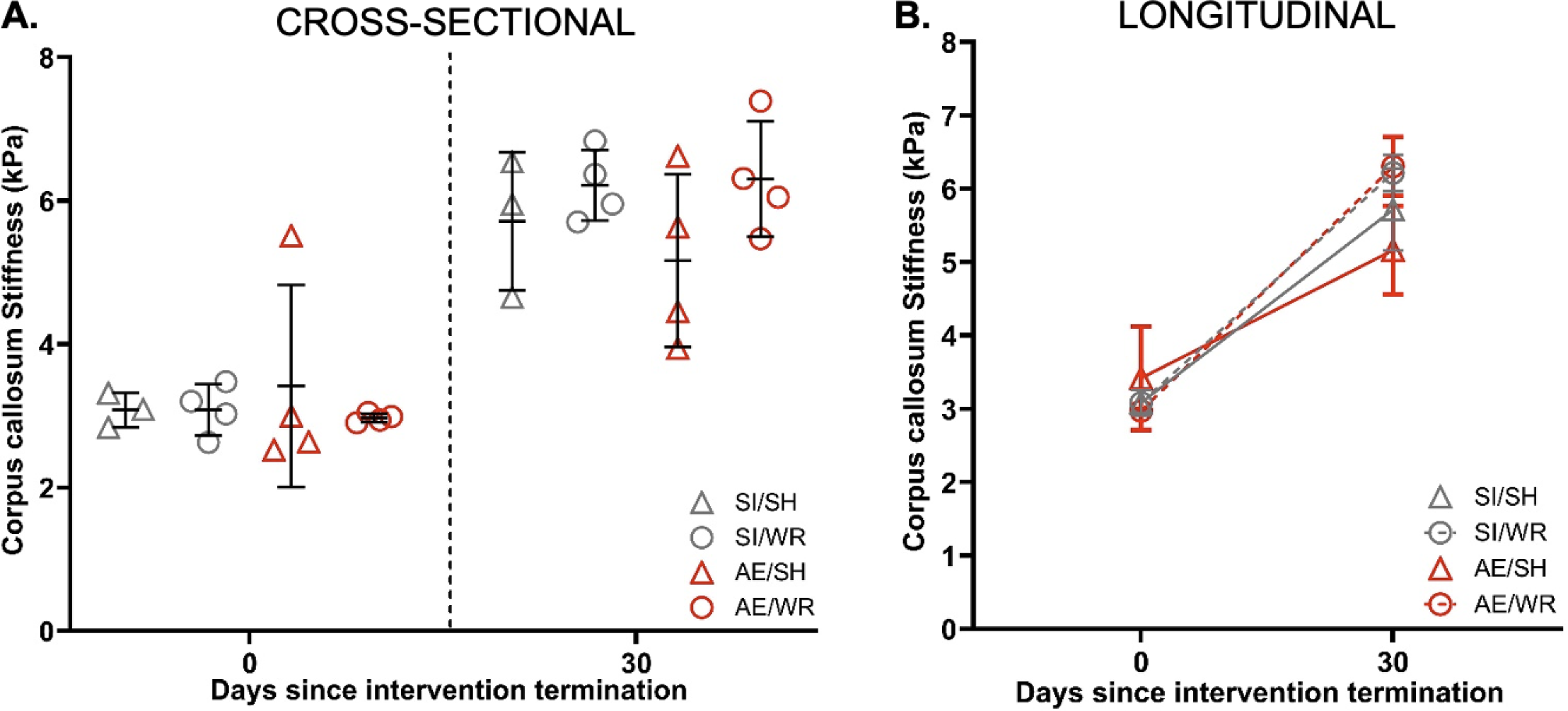
Corpus callosum shear stiffness measured immediately post-intervention and one month later in adulthood. A) Cross-sectional analysis of callosal shear stiffness data 0- and 30-days post-intervention for AE/WR, AE/SH, SI/WR and SI/SH groups. * p ≤ 0.05, error bars = SD, B) Longitudinal analysis of mean callosal shear stiffness values for AE/WR, AE/SH, SI/WR and SI/SH groups. Significant main effects and interactions not shown, error bars = ± 1 SE.

Cross sectional analysis of callosal damping ratio in 0 days post-intervention uncovered a significant main effect of postnatal treatment on damping ratio; damping ratio was lowest in AE/WR and AE/SH rats (F1 11 = 8.37, *p* = 0.01, ηp^2^ = 0.41). Similar analysis of callosal damping ratio 30 days pos-intervention did not show any significant differences between groups (**Figure 5A**). Longitudinal analysis of changes to damping ratio over time showed that callosal damping ratio increases over time in all rats (F1 11 = 66.03, *p* < 0.001, ηp^2^ = 0.86, **Figure 5B**), but that there was no significant interaction with postnatal treatment condition.

**Figure 5.**
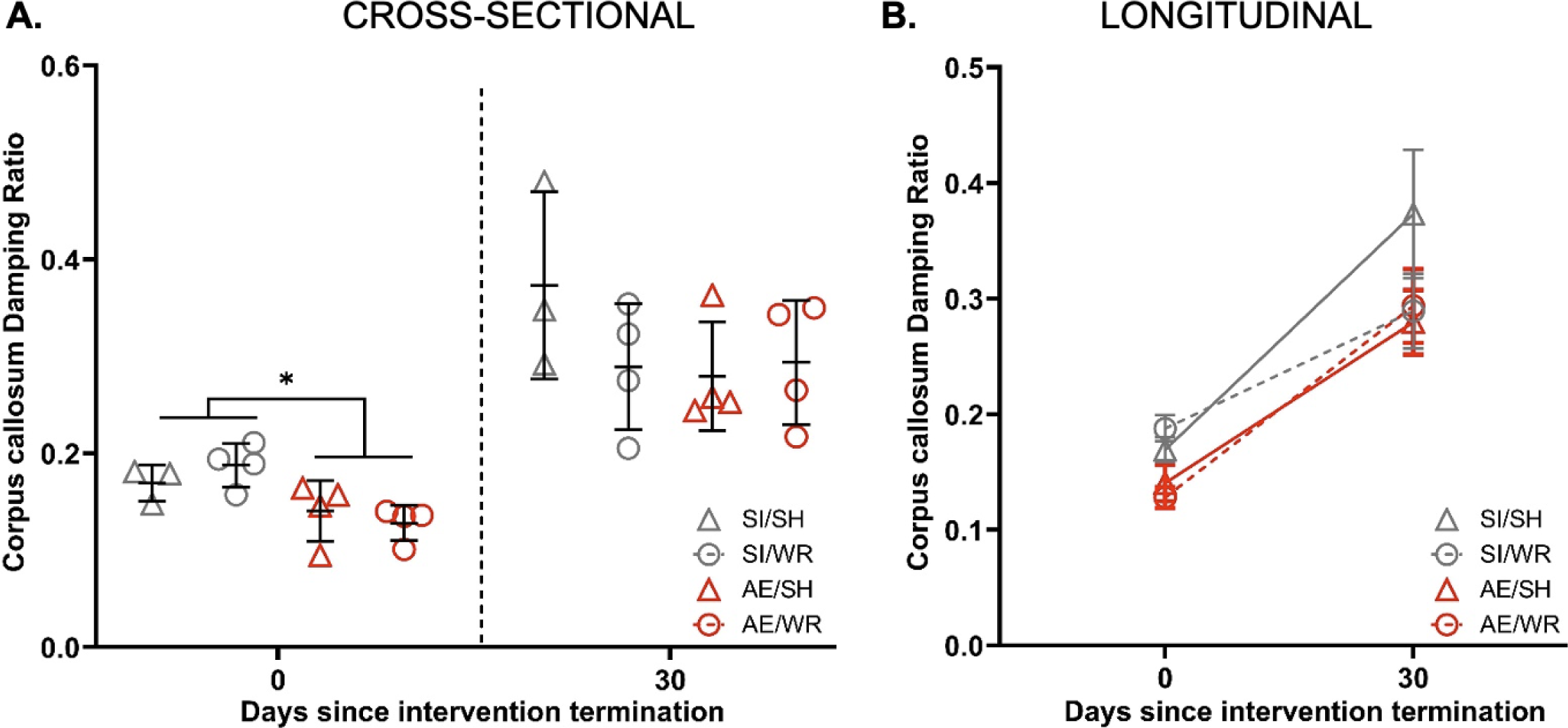
Corpus callosum damping ratio measured immediately post-intervention and one month later in adulthood. A) Cross-sectional analysis of callosal damping ratio data 0- and 30-days post-intervention for AE/WR, AE/SH, SI/WR and SI/SH groups. * p ≤ 0.05, error bars = SD, B) Longitudinal analysis of mean callosal damping ratio values for AE/WR, AE/SH, SI/WR and SI/SH groups. Significant main effects and interactions not shown, error bars = ± 1 SE. Callosal damping ratio was lowest in AE rats 0 days post-intervention, but normalized to control levels 30 days post-intervention mimicking what was observed globally.

### C. Quantification of oligodendrocytes and their precursors in the body of corpus callosum one month post-intervention

Two-way ANOVA were performed to compare the estimated number of oligodendrocytes (OLs) and oligodendrocyte precursor cells (OPCs) in the body of corpus callosum in SI/SH, SI/WR, AE/SH and AE/WR rats 30 days since intervention termination. In contrast to our original hypothesis, we discovered that adolescent exercise exposure reduced the number of OLs in AE/WR and SI/WR rats (F1 11 = 6.55, *p* = 0.03, ηp^2^ = 0.37, **Figure 6A**). The number of OPCs and the volume of the body of corpus callosum did not differ between groups in adulthood. A summary of results is described in **Table 3**.

**Figure 6.**
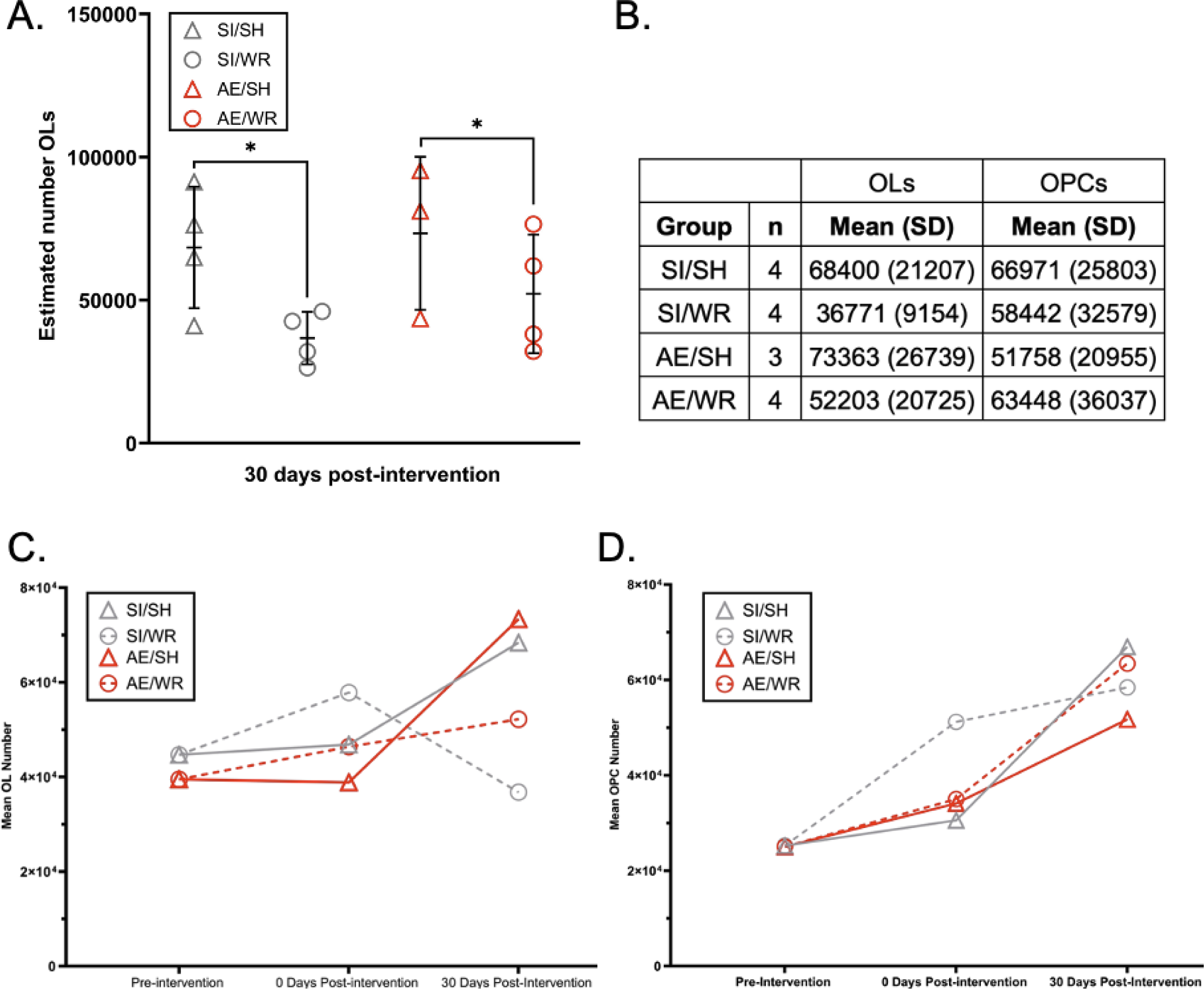
A) Cross-sectional analysis of OL number in corpus callosum 30 days post-intervention for AE/WR, AE/SH, SI/WR and SI/SH groups. * p ≤ 0.05, error bars = SD, B) Table of mean and standard deviation of OL and OPC number per group, C) Depiction of the mean OL number pre-intervention, 0 days post-intervention and 30 days post-intervention modified from Milbocker et al. (2023) and D) Depiction of the mean OPC number pre-intervention, 0 days post-intervention and 30 days post-intervention modified from Milbocker et al. (2023).

**Table 3.**
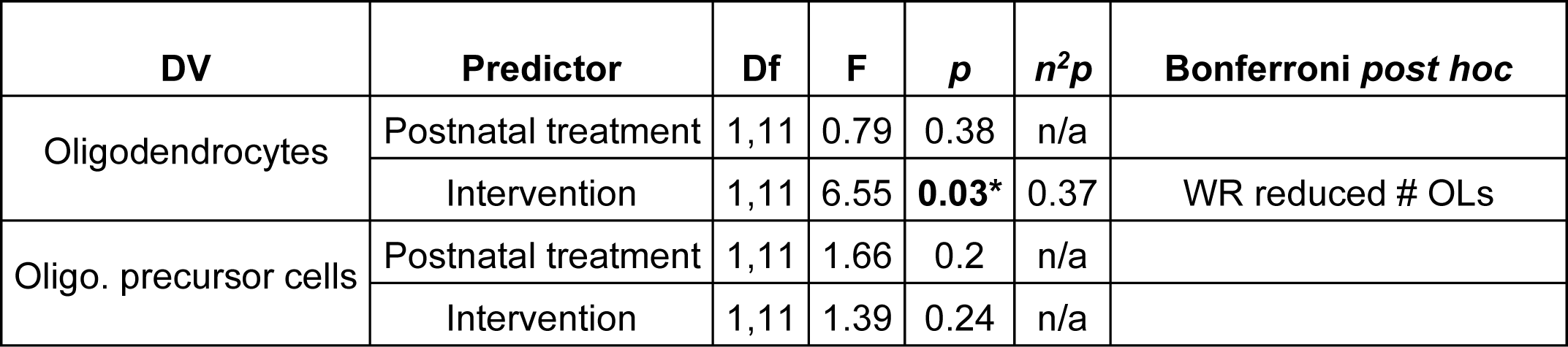
Results from factorial ANOVA comparing the effects of postnatal treatment (AE or SI control) and intervention exposure (WR or SH control) on the estimated total number of oligoglia in the body of corpus callosum 30 days DSI.

In a previous study using the same experimental paradigm, we found that mean OL and OPC number were not reduced in the AE brain pre-intervention (Milbocker et al., 2023). In that study, OPC proliferation was stimulated in the callosi of SI/WR rats 0 days post-intervention but not in the AE/WR brain (**Figure 6D**). Additionally, mean OL number was stimulated in the SI/WR and AE/WR brain 0 days post-intervention, albeit to a lesser extent in the AE brain (**Figure 6C**).

Independent samples t-tests run within each treatment (AE or SI) and intervention (SH or WR) group revealed that the number of OLs did not differ from 0 to 30 days post-intervention in SI/SH rats (*p* = 0.15, **Figure 6C**). However, the number of OPCs increased from 0 to 30 days post-intervention in this group (*p* = 0.01, **Figure 6D**). In contrast, the number of OPCs did not change between 0 to 30 days post-intervention in AE/SH rats (*p* = 0.05, **Figure 6D**), while the number of OLs increased from 0 to 30 days post-intervention in this group (*p* = 0.01, **Figure 6C**). Importantly, the OL number decreased from 0 to 30 days post-intervention in SI/WR rats (*p* = 0.01, **Figure 6C**) and there was no change in the number of OPCs (**Figure 6D**). There was no significant change in OL number in the AE/WR rats from 0 to 30 days post-intervention (**Figure 6C**), yet the number of OPCs was significantly increased between these time points (*p* = 0.03, **Figure 6D**).

## DISCUSSION

This study examined the effects of an adolescent exercise intervention on forebrain and corpus callosum development in a rat model of FASD with alcohol exposure during the brain growth spurt. For the first time, noninvasive MRE scanning was used to assess alterations to the mechanical properties (stiffness and damping ratio) of forebrain and corpus callosum in FASD model. Synthesizing our findings, we observed that AE appears to result in lower stiffness (AE/SH group) and damping ratio (both AE/SH and AE/WR groups) of the brain in adolescence. Lower brain stiffness is consistent with reduced brain tissue integrity as expected from previous studies of AE at a similar developmental time point and is consistent with lower stiffness due to neurological conditions in many human MRE studies on adults (Murphy et al., 2019). The developmental brain MRE literature is only just emerging, but a study on human children and adolescents with cerebral palsy also found lower brain stiffness (Chaze et al., 2019). On the other hand, a rat brain MRE study found no differences in stiffness in poly (I:C)-induced maternal immune-activated rats (Liu et al., 2023), which to our knowledge is the only other rat brain MRE study of a neurodevelopmental disorder. There are no reports on brain mechanical properties assessed with MRE in clinical or preclinical studies of FASD.

Our study was designed to consider immediate and persistent effects of aerobic exercise on brain mechanical properties in FASD. From cross-sectional analysis of data immediately after intervention, where brain stiffness was lower only in AE/SH rats but not in AE/WR rats, we conclude a positive effect of exercise on brain tissue integrity. Longitudinal analysis further revealed that forebrain stiffening occurred in AE/SH rats from 0 to 30 days post-intervention, effectively matching the stiffness of the other groups. These data support our hypothesis that adolescent exercise intervention aids the developmental stiffening of the forebrain in FASD rats, mitigating alcohol-induced alterations to neurodevelopmental trajectory. Moreover, these findings are in line with the clinical results by Schwarb et al. (2017) and Sandroff et al. (2017) that demonstrated that MRE properties reflect positive brain health and tissue integrity following aerobic activity. It is postulated that a combination of alterations to vasculogenesis, brain perfusion, neurogenesis, and synaptogenesis, known to occur with exercise (Klintsova et al., 2013; Voss et al., 2013), may contribute to the observed increase in brain stiffness after exercise. These results suggest exercise could be pivotal for reducing disparities in learning deficits, legal issues, and behavioral impairments in youth with FASD.

We also found that the damping ratio of the forebrain and corpus callosum was lower in AE rats compared to SI rats in adolescence immediately post-intervention. Damping ratio is a biomechanical property of viscoelastic tissue that describes the relative viscous-to-elastic behavior of the material and determines energy dissipation and motion attenuation. In MRE, damping ratio and other parameters that describe similar viscoelastic behaviors are considered to reflect microstructural tissue organization (Sack et al., 2013). Given these principles, we suspect that the observed reduction to forebrain damping ratio in AE rats in adolescence may reflect delayed axonal pruning that is characteristic of the FASD adolescent brain (Fernández-Jaén et al., 2011; Gursky et al., 2020; Smith et al., 2022b; Treit et al., 2014; Yang et al., 2012). The excess of axonal connections that persists in late adolescence in the FASD brain could be lowering energy dissipation. The increase in damping ratio in AE rats over time to reach the levels of the SI control rats further suggests a developmental delay captured by damping ratio. Though we note there appeared to be no intervention effect on damping ratio such that it is unaffected by exercise, suggesting a possible different microstructural mechanism driving this parameter compared to stiffness.

Our longitudinal MRE measures are also confounded by normally occurring changes in brain tissue mechanical properties with development. This is most clearly observed in measures of the corpus callosum, where all groups increased in stiffness and damping ratio between adolescence and adulthood, with differences in degree of change occurring based on postnatal treatment or intervention condition. The effects of aging on brain mechanical properties in adulthood are well-known (Hiscox et al., 2021), with lower stiffness (Arani et al., 2015; Hiscox et al., 2018; Sack et al., 2011) and higher damping ratio (Delgorio et al., 2021) occurring with older age. Much less is known about periods of brain development and maturation. Several studies have reported global stiffness to be generally consistent with age in children, adolescents, and young adults (McIlvain et al., 2018; Ozkaya et al., 2021; Yeung et al., 2019), though a recent study on a large sample found instead lower stiffness and higher damping ratio with increasing age from 5 to 35 years (McIlvain et al., 2022). This is in contrast to rodent brain MRE studies that have reported increased stiffness and either decreased or unchanged phase angle (comparable to damping ratio) with age (Guo 2019, Schregel 2012), though with regional differences in both magnitude and direction of age-related property change.

The disagreement between only a small number of studies on how brain mechanical properties are affected by age in younger groups, coupled with the seemingly different age-related trends between humans and rodents, makes interpreting the damping ratio results in our study more challenging. Damping ratio is less commonly reported than stiffness in part due to methodological challenges and differences between MRE protocols. Using the same NLI methods as in this study, we have previously observed in healthy adult humans that lower damping ratio was associated with greater performance on cognitive tasks (Johnson and Telzer, 2018; Schwarb et al., 2017), suggesting that lower damping ratio indicates greater tissue integrity. We also observed higher damping ratio in brain tissue of children with cerebral palsy (Chaze et al., 2019). Our finding of reduced damping ratio in FASD appears inconsistent with these findings, but this may be due to different aging effects in younger groups or rodents, as discussed above, or a different manifestation of pathology not well captured by the limited number of comparable studies that currently exist. As the field continues to grow more studies will illuminate the mechanistic interpretation of our results, including future examinations of humans with FASD.

Towards this effort, we quantified the number of precursor and myelinating oligoglia in corpus callosum one-month post-intervention in all rats from this study. It is predicted that the integrity of the oligodendrocyte matrix contributes to the mechanical properties of white matter. Indeed, global and regional stiffness is reduced (Millward et al., 2015) and loss modulus, a similar measure to damping ratio, decreased (Schregel et al., 2012) in rodent models of demyelinating diseases like multiple sclerosis where oligodendrocyte apoptosis occurs. Several studies on rodent and non-human primate models of FASD have uncovered that AE reduces the number of OPCs and OLs in the adolescent brain (Creeley et al., 2013; Milbocker et al., 2023; Newville et al., 2017). Similar reductions to oligoglia population were observed in fetal brain tissue with known prenatal alcohol exposure (Darbinian et al., 2021). Persistent hypomyelination was identified via quantification of the g-ratio on electron micrographs in the posterior corpus callosum of mice exposed to alcohol during the brain growth spurt (Newville et al., 2021). Moreover, voluntary wheel running has been shown to attenuate loss of OLs in demyelination disease (Mandolesi et al., 2019). Thus, we expected that MRE would be an advantageous tool for identifying microstructural changes to the rodent brain following exercise intervention in a rat model of FASD where developmental myelination is perturbed. In a previous study, we showed that voluntary aerobic exercise in adolescence increased the number of OPCs in the body of corpus callosi of SI rats only (Milbocker et al., 2023). In addition, we found that the number of mature OLs was increased in both SI/WR and AE/WR rats 0 days post-intervention, suggesting that moderate myelin plasticity was achieved by exercise intervention exposure in the FASD brain (i.e., oligodendrocyte maturation without precursor cell proliferation). In the current study, we expanded on these findings by quantifying OPCs and OLs in the same region one month post-intervention in early adulthood. Our results indicate that the volume of the body of corpus callosum and the population of OPCs did not differ significantly between groups 30 days post-intervention. This suggests that OPC proliferation was halted following exercise cessation in SI rats. Unexpectedly, OL number was reduced in both AE/WR and SI/WR adult rats 30 days post-intervention, suggesting a negative rebound in OL survival in white matter following detraining. While this phenomenon has yet to be discovered in respect to glia of the CNS, there is ample evidence that adolescent voluntary exercise cessation leads to a negative rebound in adult neurogenesis in rodents (Hopkins et al., 2011; Nishijima et al., 2017). A negative rebound describes the deleterious effect of the cessation of an external stimulus on internal processes below control (or starting) levels. Thus, it would be inappropriate to postulate that voluntary exercise cessation led to a negative rebound on OPC proliferation as OPC quantity in rats exposed to the adolescent intervention was not below that of controls in adulthood. Importantly, this effect was not mirrored by results from noninvasive MRE scanning, indicating a current limitation of our protocol to perform “virtual histology.”

There are two mechanisms by which exercise cessation might lead to a reduction in the number of OLs in the corpus callosum below control levels. First, exercise cessation may arrest the maturation of pre-myelinating OLs (preOLs) to mature OLs via genetic mutation of mTOR which increases the susceptibility of preOLs to Bax-mediated apoptosis (Hughes and Stockton, 2021; Ornelas et al., 2020). Indeed, voluntary exercise upregulates mTOR levels in glia of the CNS in rats (Lloyd et al., 2017), and thus exercise cessation may cause a reduction to mTOR production. Second, exercise cessation may cause metabolic distress in preOLs. Oxidative stress resulting from mitochondrial dysfunction prevents the maturation of preOLs to myelinating OLs, and thus alterations to oligoglia bioenergetics may increase the susceptibility of preOLs to apoptosis (Harris and Attwell, 2012; Rinholm et al., 2011; Yan and Rivkees, 2006). Finally, rats with voluntary exercise show a reduction to p53 in several gray matter regions (Rezaee et al., 2019), and p53 deficiency has been shown to benefit remyelination of the corpus callosum in rodent models of multiple sclerosis and stroke (Luo et al., 2021). Thus, increased levels of p53 resulting from exercise cessation could induce OL cytotoxicity, temporarily reducing the number of OLs below control levels in the brain. Importantly, in studies documenting a negative rebound effect of exercise cessation on neurogenesis in rats, this effect was found to be transient and neurogenesis was returned to control levels approximately 2 months following exercise termination (Nishjima et al., 2017).

There are several limitations that restrict the interpretation of MRE and histological findings in our study. First, we did not capture MRE scans at a pre-intervention time point which limits our interpretation of how exercise intervention specifically impacts forebrain and callosal stiffness and damping ratio in AE and SI rats. Second, this pilot study had a low sample size. Albeit the results demonstrate that AE does impact brain stiffness and damping ratio in adolescence. Third, improvements to NLI and the MRE setup and acquisition are in progress to increase the feasibility of subregional property estimation. Given the findings by Newville et al. (2017, 2021), it is possible that greater differences in the mechanical properties of the posterior corpus callosum might be observed following intervention exposure. Finally, adding an additional scanning time point between PD42 and PD70 may have also revealed exercise-related effects as there would be a lower contribution from rodent age.

## CONCLUSION & CLINICAL RELEVANCE

In summary, this pilot study was the first to investigate the effects of alcohol exposure and exercise intervention on brain mechanical properties, *in vivo*, in a rodent model of FASD. Properties were estimated using MRE and nonlinear inversion with similar quality to human studies, making these preclinical results clinically translatable. Ultimately, our findings from both MRE and histology strongly support that exercise improves development in FASD rats. Delayed development due to FASDs is currently untreatable in humans, but exercise may help them catch up to baseline. As MRE is a noninvasive, indirect measure of tissue integrity, it can be used to monitor progress. Thus, there is great promise for MRE as an evaluation tool for behavioral therapies.

## Data Availability Statement

The data that support the findings of this study are available on request from the corresponding author. The data are not publicly available due to privacy or ethical restrictions.

